# ClonoScreen3D-CRISPRi Uncovers Genetic Modifiers of Radiation Response in Glioblastoma

**DOI:** 10.64898/2026.04.17.719014

**Authors:** Sukjun Lee, Anke Husmann, Jianbin Li, Chao Zheng Li, Sunil Modi, Saif Ahmad, Simon MacKay, Andrew Paul, Mark R Jackson, Anthony J Chalmers, Nicola McCarthy, Natividad Gomez-Roman, Erica Bello

## Abstract

**Background:** Glioblastoma (GBM) is the most aggressive primary brain tumor in adults. Radioresistance, partly mediated by glioma stem-like cells, represents a major clinical challenge which could be overcome by the identification of the modulators of radioresistance. Existing CRISPR screens in human GBM models have largely used two-dimensional cultures with short-term viability readouts, failing to capture the long-term clonogenic behaviour underlying tumour recurrence after radiotherapy.

**Method:** We developed ClonoScreen3D-CRISPRi, combining CRISPRi-mediated gene knockdown with three-dimensional clonogenic survival assays. Two GBM cell lines (G7 and GBML20), differing in MGMT promoter methylation status, were engineered to express the KRAB-dCas9 editor. Nine candidate radiosensitivity modifiers, selected through transcriptomic analysis, pharmacological studies, and literature review, were examined in both lines. Target validation was performed using full radiation dose–response assays and a pharmacological inhibitor.

**Results:** The majority of candidate genes significantly altered survival fraction following irradiation in both cell lines. Knockdown of NFKB2, RELB, and CDK9 produced the most potent radiosensitization, with sensitizer enhancement ratios of 1.39–1.70 in validation studies — exceeding those of established radiosensitizers including PARP and ATM inhibitors. Notably, knockdown of these genes induced no significant cytotoxicity in the absence of radiation. Pharmacological validation using an IKKα inhibitor confirmed these findings, implicating non-canonical NF-κB signalling and CDK9-dependent transcriptional elongation as critical adaptive mechanisms in GBM radioresistance.

**Conclusions:** ClonoScreen3D-CRISPRi is a scalable, physiologically relevant platform for identifying genetic modifiers of radioresistance. The non-canonical NF-κB pathway and CDK9 represent promising radiosensitizing targets, and larger screens could enable systematic prioritisation of candidates for clinical translation.

**Key Points:**

- ClonoScreen3D-CRISPRi combines gene knockdown with 3D clonogenic survival assays
- We identified *NFKB2, RELB*, and *CDK9* as modifiers of radioresistance in two GBM cell lines
- Validation experiments show ClonoScreen3D-CRISPRi reliably identifies radiosensitizers in GBM

**Importance of the study:** Glioblastoma (GBM) remains one of the most lethal human cancers, with radioresistance representing a central barrier to improved patient outcomes. While CRISPR-based screens have begun to illuminate genetic drivers of GBM biology, prior approaches using human models have largely relied on two-dimensional culture systems and short-term viability readouts that inadequately model the disease. This study introduces ClonoScreen3D-CRISPRi, a novel platform that integrates CRISPRi-mediated gene knockdown with three-dimensional clonogenic survival assays in patient-derived GBM cells — more faithfully recapitulating the long-term clonogenic potential that underlies post-radiotherapy recurrence. Using this platform, we identified *NFKB2, RELB*, and *CDK9* as potent genetic modifiers of radioresistance, with sensitizer enhancement ratios exceeding those of established clinical radiosensitizers such as PARP and ATM inhibitors. Pharmacological validation of the non-canonical NF-κB pathway demonstrates direct translational relevance, providing a rationale for targeting this axis in combination with radiotherapy to improve GBM treatment.

## Introduction

Glioblastoma (GBM) remains the most aggressive and lethal primary brain tumour in adults, characterized by extensive heterogeneity, rapid recurrence, and poor prognosis despite multimodal treatment^1,2^. Standard-of-care therapy comprises neurosurgical resection followed by radiotherapy and temozolomide chemotherapy, however this extends median survival to only ∼15 months^1^. This limited efficacy reflects intrinsic and acquired radioresistance in GBM often driving tumour relapse^3^, which poses major clinical challenges.

A contributing factor to radioresistance is the presence of glioma stem-like cells (GSCs), a subpopulation capable of self-renewal and differentiation. These cells exhibit constitutive activation of DNA damage response and cell cycle checkpoint pathways, enabling efficient survival under genotoxic stress. This is characterized by persistent activation of damage-sensing and checkpoint kinases, including Chk1, Chk2, ATR, ATM, RAD17, and RAD51, accompanied by upregulation of key DNA repair enzymes such as PARP1 and TIE2 ^4–6^. This heightened repair capacity underpins the intrinsic radioresistance of GBM and promotes tumour recurrence. Therefore, identifying modulators of GBM radioresistance holds critical importance for developing effective radiosensitization therapies with the aim of improving clinical outcomes.

Recent CRISPR-Cas9-based functional genomics approaches in human GBM models have enabled systematic identification of genetic drivers of GBM^7–11^. These studies have highlighted roles for DNA repair, chromatin remodelling, and cell cycle regulators in GBM, suggesting their potential as prospective therapeutic targets. However, most studies were conducted in 2D monolayer cultures, which fail to recapitulate the complex microenvironment of GBM. In addition, many rely on short-term viability assays, providing limited insight into long-term clonogenic capacity, a key determinant of tumour recurrence following radiotherapy. Therefore, genetic screens in 3D complex human models of GBM are needed to maximize the chances of uncovering novel targets for the effective treatment.

To overcome the limitations of current screening approaches in GBM, we have made use of ClonoScreen3D, a previously developed 3D clonogenic screening platform in 96-well plate format which effectively recapitulates the key clinical features of GBM as well as responses to radiation and molecular therapies ^12^. ClonoScreen3D allows screening of novel radiation-drug combinations using clonogenic survival as readout. In this study, we optimized and adapted this platform for genetic screening, incorporating transient CRISPRi-mediated gene knockdown (KD) with irradiation treatment to enable the assessment of gene function in radioresistance. We demonstrated that this approach can identify genetic modifiers of radioresistance in GBM and that a scalable workflow to enable larger CRISPRi screens is feasible.

## Methods and Materials

### Cell culture

Patient-derived G7 cells were obtained from Professor Colin Watts, as previously described^13^ Gene mutations commonly observed in GBM and present in these lines have been previously published^12^. Patient-derived GBML20 GBM cells were obtained from Dr. Dimitris Placantonakis (NYU). The GBM cell lines G7 and GBML20 were cultured as monolayers on Matrigel (Merck)-coated plates (1:40 dilution in basal medium without supplements) using serum-free, cancer stem cell-enriching medium. G7 cells were maintained in Advanced DMEM/F12 (Thermo Fisher Scientific) supplemented with 1% B27, 0.5% N2 (both from Thermo Fisher Scientific), 5 µg/mL heparin, 10 ng/mL fibroblast growth factor 2 (Merck), 20 ng/mL epidermal growth factor (Merck), 1% GlutaMAX, and 1% penicillin–streptomycin (Thermo Fisher Scientific). GBML20 cells were maintained in Neurobasal™ Medium (Thermo Fisher Scientific) supplemented with 2% B27 (minus Vitamin A), 1% N2, 1% non-essential amino acids, 1% GlutaMAX, and 1% penicillin–streptomycin (Thermo Fisher Scientific). The CRISPRi expressing versions of these lines (see below) were maintained in the same culture medium used for their wild-type counterparts. The confluency of all GBM cell lines was maintained below 70%, and only cultures within 10 passages after thawing from frozen stocks were used throughout this study. For virus production, HEK293FT cells were cultured in high-glucose DMEM (Thermo Fisher Scientific) supplemented with 10% FBS, 1% non-essential amino acids, 1% GlutaMAX, and 1% penicillin–streptomycin (Thermo Fisher Scientific).

### CRISPR interference (CRISPRi) engineering of GBM cell lines

G7 and GBML20 cell lines were engineered for constitutive expression of the Zim3 KRAB domain fused to dead Cas9 (dCas9)^14^ via lentiviral transduction. Lentiviral vectors encoding Zim3-dCas9-P2A-mCherry (Addgene #188766) were produced in H293FT cells using a third-generation packaging system. Briefly, H293FT were co-transfected with Zim3-dCas9, psPAX2 (Addgene #12559), and pMD2.G (Addgene #12260), after which viral supernatants were collected and concentrated with Lenti-X, subsequently aliquoted for storage at –80 °C. To generate stably expressing Zim3-dCas9 GBM lines, monolayer GBM cultures at approximately 70% confluence were spin-infected with polybrene (8 ug/ml) and the thawed packaged Zim3-dCas9 lentivirus. After one passage, the top 20% of mCherry-expressing cells was collected by flow cytometry, and the procedure was repeated one week later to further enrich for high expressors. These sorted populations were used for all subsequent experiments.

### Reverse-transfection of single-guide RNAs (sgRNA) for CRISPRi-mediated gene KD

Reverse transfection of sgRNAs was performed using Lipofectamine RNAiMAX (Thermo Fisher Scientific) in accordance with the manufacturer’s instructions. Briefly, sgRNAs were diluted to a final concentration of 0.6 µM in 80 µL Opti-MEM (Thermo Fisher Scientific), and combined with an equal volume of Opti-MEM containing 1.3 µL RNAiMAX. Following a 20-minute incubation at room temperature for complex formation, the transfection mixture was added to 16,000 of CRISPRi effector–expressing GBM cells suspended in 160 µL Opti-MEM supplemented with 10% FBS and seeded onto Matrigel-coated 48-well plates. These cells were maintained for 72 hours for CRISPRi-mediated KD, without media replacement. For dual-sgRNA transfections, both sgRNA were prepared in the same protocol at a final concentration of 0.6 µM each, and RNAiMAX was increased proportionally to 2.6 µL. After 72 hours of transfection, cells were detached using StemPro Accutase (Thermo Fisher Scientific), and optimal number of viable cells were transferred to Matrigel-coated Alvetex plates (Reprocell). The remaining cells were stored in TRI Reagent (Zymo Research) at −80 °C for subsequent qPCR analysis to verify KD. After 24 hours of seeding, cells were subjected to irradiation or compound treatment, as described in detail in the following section. sgRNAs were either designed based on sequences reported in Replogle et al^14^. or using CRISPick (Broad Institute, https://portals.broadinstitute.org/gppx/crispick/public)^15,16^, and synthesized by Synthego.

### ClonoScreen3D clonogenic survival assay

ClonoScreen3D assays were performed as previously described using 96-well Alvetex plates^12^. For 3D Alvetex cultures, plates were first treated with 70% ethanol for membrane hydration, followed by washing twice with PBS. After washing, plates were coated with Matrigel coating media (1:40 dilution in basal medium without supplements, as used for cell culture). Optimal seeding densities and radiation doses for each GBM cell line in the 96-well assay were determined through preliminary optimisation studies; 160 cells per well were used for G7/G7-Zim3 and 180 cells per well for GBML20/GBML20-Zim3, while radiation doses of 3 Gy were selected for both cell lines. Cells were seeded onto Alvetex plates and irradiated 24 hours after seeding using an RS-1800Q Biological Irradiator (Rad Source Technologies, USA). Irradiation was performed using 160 kV X-rays at a dose rate of 0.85 Gy/min (25% power setting). For compound treatments, cells on Alvetex plates were exposed to the indicated compounds 2 hours before irradiation. Details of all compounds used in this study are described in Supplementary Table S2. Following irradiation, plates were incubated at 37 °C with 5% CO_2_for 14 days, after which colonies were stained with thiazolyl blue tetrazolium bromide (MTT) and fixed with 2% paraformaldehyde (PFA). Images were captured using a Syngene G:Box imaging system for subsequent colony counting analysis.

### Full radiation dose response

A full radiation dose–response assay using 12-well Alvetex plates was performed to validate the radiosensitizing effects of CRISPRi-mediated gene KD. Plates were prepared using the same Matrigel coating procedure described above. sgRNA transfection was performed in 48 well plates as previously detailed, and cells were detached 72 hours post-transfection before being seeded according to the intended radiation dose. For radiation, 500 cells per well were seeded for 0, 1, 2, and 3 Gy, and 800 cells per well for 4 and 5 Gy. Irradiation was performed 24 hours after seeding, and colony staining, image acquisition, and subsequent analysis were conducted as described above, 14 days after irradiation. Sensitizer enhancement ratios (SERs) were calculated from mean inactivation doses derived from linear–quadratic model fits (https://jackson-chalmers.shinyapps.io/lq_app/).

### Image Analysis Pipeline of colony counting

The image analysis pipeline is implemented in three sequential parts: initial image segmentation with Cellpose (v3.1.1.1 using cyto3 model, https://www.cellpose.org) to find wells^17^, followed by a pipeline within CellProfiler (v4.2.6, https://cellprofiler.org) to crop the image of the plate into individual images of wells^18^, and subsequent quantitative analysis using CellProfiler to count colonies. Raw images are first processed with Cellpose, a highly specialized open-source software for segmentation of biological objects, to segment wells, by utilising the fact that the wells are round objects. Cyto3 model is based on the Segment Anything Model which had been further trained on cells. Setting the diameter to the diameter of the wells, the model creates masks for the wells. The original image with the CellPose masks is further processed in CellProfiler, an open-source software for high-throughput biological image analysis, to determine a grid of objects (wells) along which lines the original image is cropped into a set of individual images of wells. The number of columns and rows was adjusted by hand. The cropped well images are analysed in a second pipeline in CellProfiler. Each well image is cropped circularly to exclude shadows and artifacts outside the well area. Illumination corrections, background subtraction and noise reduction improve segmentation accuracy. Colonies are segmented using the Otsu thresholding method (two-class mode). Thresholds are adjustable to optimize segmentation. Shape-based criteria are applied to ensure accurate identification of round colonies. Quantitative measurements are extracted for each colony, including size, shape descriptors, and intensity features. The pipeline produces annotated images showing detected maxima and outlines of segmented colonies overlaid on the original images. Additionally, a .csv file containing size, shape, and intensity metrics for each colony is exported for downstream analysis.

### Flow Cytometry analysis

To assess CRISPRi activity, cells transfected with sgRNAs targeting specific genes were analysed by flow cytometry. After 96 hours of transfection, cells were detached using Accutase and stained with the LIVE/DEAD Fixable Dead Cell Stain Kit (Thermo Fisher Scientific) for 30 minutes. Cells were then fixed and permeabilized using the eBioscience Intracellular Fixation & Permeabilization Buffer Set for 30 minutes, followed by one wash with eBioscience Flow Cytometry Staining Buffer (Thermo Fisher Scientific). Subsequently, cells were incubated with primary antibodies for 1 hour at 4 °C under the dark conditions described in Supplementary Table 3. After staining, cells were washed three times with 500 µL of Flow Cytometry Staining Buffer and resuspended in the same buffer for acquisition on a CytoFLEX flow cytometer (Beckman Coulter).

### qPCR analysis

Total RNA of each transfected cell was extracted using the Direct-zol RNA Microprep Kit (Zymo Research). cDNA was synthesized from 0.5 µg of total RNA of each transfected cells using the Transcriptor First Strand cDNA Synthesis Kit (Roche). qPCR was performed using SYBR Green I Master Mix (Thermo Fisher Scientific) on a QuantStudio 7 PRO PCR System (Thermo Fisher Scientific). The primers used in this study are listed in Supplementary Table S4.

### Proliferation assay

Cell growth was evaluated using the RealTime-Glo MT Cell Viability Assay (Promega). A total of 1,000 cells were seeded per well in Matrigel-coated (1:40 dilution, as used for cell culture) white 96-well luminescence plates, with all conditions plated in triplicate. Immediately after seeding, 2X RealTime-Glo reagent containing both substrate and enzyme was added to each well. After a 30-minute incubation, luminescence was measured to obtain the baseline signal (day 0), and subsequent measurements were recorded every 24 hours for 6 days. Luminescence values at each time point were normalized to the corresponding day 0 measurement.

### Extreme limiting dilution assay (ELDA)

To quantify cancer stem cell frequency, cells were seeded at decreasing densities (80, 40, 20, 10, and 5 cells per well) into 96-well flat-bottom plates containing serum-free, cancer stem cell-enriching medium. Each cell density was plated in six technical replicates per biological repeat, with three biological repeats performed in total. Cultures were maintained for 21 days, with medium replenished on days 8 and 15. At the endpoint, cells were fixed with 4% formaldehyde, and spheres larger than 20 µm in diameter were counted per well using a phase-contrast microscope. Sphere-forming frequency was calculated using the ELDA software (http://bioinf.wehi.edu.au/software/elda/) with the default confidence interval of 0.95. Data analysis performed likelihood ratio tests in accordance with a generalized linear model^19^.

### Protein Expression Analysis

Protein expression levels in GBM tumours relative to normal healthy brain tissue was performed using the CPTAC (Clinical Proteomic Tumour Analysis Consortium) database (https://cptac-data-portal.georgetown.edu/)^20^ on tissues using isobaric tandem mass tags (TMT). For this study, our analysis was restricted to GBM brain cancer cohorts to ensure pathological relevance. The integration and processing of these proteomic datasets were performed as previously described^20^. Briefly, raw protein expression values were retrieved from the CPTAC data portal and subjected to log_2_normalization within each sample. To enable standardized comparisons across the tumour and control cohorts, a Z-score was calculated for each protein, defined as the number of standard deviations from the median across all samples. These normalized values were utilized to compare protein abundance in GBM tissues relative to healthy brain controls. Statistical significance was evaluated using two-sample unpaired t-tests.

### Statistical analysis

To evaluate CRISPRi-mediated KD, qPCR (ΔCt) and FACS (MFI) data were analysed using paired two-tailed t-tests. For comparisons involving multiple groups, one-way ANOVA or repeated-measures ANOVA (RM-ANOVA), followed by Dunnett’s or Tukey’s multiple comparisons test, was applied as appropriate. Plating efficiency (PE) and surviving fraction (SF) from ClonoScreen3D-CRISPRi screens were analysed using the same ANOVA–Dunnett framework. Pearson correlation analysis was used to assess concordance between screening datasets, and between automated and manual colony counting. All statistical analyses were performed using GraphPad Prism version 11.

## Results

### Generation of CRISPRi-expressing GBM lines

To develop a method to identify genetic modifiers of radiation response in a 3D model of GBM, we integrated CRISPRi-mediated gene KD in GBM patient-derived cell lines within the ClonoScreen3D platform. We selected two GBM patient-derived lines, G7 and GBML20, that differ in the methylation status of the O^6^-methylguanine-DNA methyltransferase (*MGMT*) promoter - methylated in G7 and unmethylated in GBML20. This molecular feature influences DNA repair capacity, cellular responses to chemotherapy, clinical responses to treatment and patient prognosis and is used clinically to guide treatment decisions^2^. We engineered G7 and GBML20 cells to constitutively express the KRAB Zim3-dCas9 CRISPRi system^5^ and confirmed that expression of Zim3-dCas9 was not silenced after repeat passaging of the cells (Supplementary Figure 1A). Moreover, Zim3-dCas9 expression did not affect cell growth (Supplementary Figure 1B), response to radiation (Figure 1A) or clonogenic potential (Figure 1B). The IC_50_ response to the known radiosensitizer olaparib was unaffected, indicating that the engineered cell lines retain key features of their GBM wild-type counterparts (Figure 1C).

**Figure 1.**
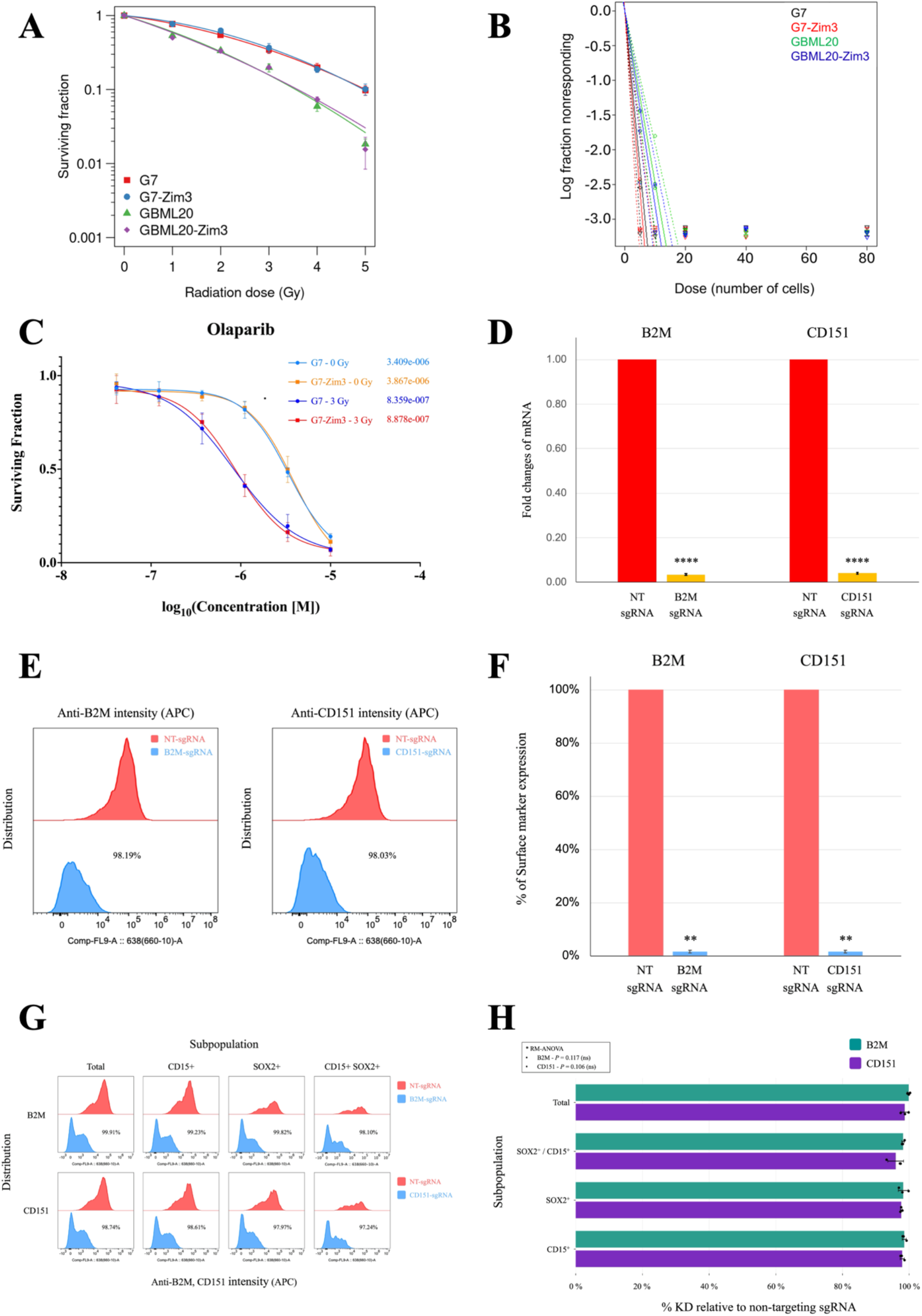
Establishing efficient gene knockdown in GBM lines by CRISPRi. (A) Full Radiation Dose response curves of parental and CRISPRi-engineered GBM cell lines. Surviving fraction was assessed following increasing doses of ionizing radiation in G7, G7-Zim3, GBML20, and GBML20-Zim3 cells, showing comparable baseline radiosensitivity between parental and Zim3-expressing lines. Data fitted using the linear quadratic model. (B) Extreme limiting dilution neurosphere assay of parental and CRISPRi GBM lines. Log-transformed fraction of non-responding wells is plotted against the number of cells plated per well, indicating similar sphere-forming capacity between parental and Zim3-expressing cells in both G7 and GBML20 backgrounds. (C) Validation of drug response in G7-Zim3 cells using olaparib. Clonogenic survival of G7 glioblastoma cells following treatment with olaparib or DMSO administered 2 h prior to 3 Gy irradiation. The surviving fraction (SF) was calculated relative to the corresponding DMSO-treated irradiated control to normalize. IC_50_ values under both non-irradiated (0 Gy) and irradiated (3 Gy) conditions are shown for parental G7 and G7-Zim3 cells. Data represent mean ± SD from independent biological replicates. (D) Quantitative PCR analysis validating CRISPRi-mediated KD of gene. KD efficiency of *B2M* and *CD151* was quantified independently by qPCR and expressed as relative mRNA levels normalized to matched NT-sgRNA controls. Statistical significance for each gene was assessed using paired two-tailed t-tests performed on ΔCt values from biological replicates (n = 3). Data are presented as mean ± SD, \*\*\*\**P* < .0001. (E) Flow cytometry histograms showing surface protein knockdown by CRISPRi. Representative flow cytometry histograms demonstrating robust knockdown of B2M and CD151 surface expression following sgRNA-mediated targeting compared with NT-sgRNA control cells. The marked leftward shift in fluorescence intensity indicates efficient depletion of the respective cell surface proteins. (F) Quantification of surface marker knockdown efficiency. The percentage reduction in surface marker expression for B2M and CD151 following CRISPRi-mediated knockdown was determined by flow cytometric analysis based on isotype-corrected mean fluorescence intensity (MFI), relative to non-targeting sgRNA controls. Statistical significance between NT-sgRNA and gene-specific sgRNA conditions was assessed using paired two-tailed t-tests across biological replicates (n=3). Data are presented as mean ± SD, ** *P* < .01 (G) Representative flow cytometry histograms of B2M and CD151 surface expression across indicated subpopulations following CRISPRi-mediated knockdown. Red traces represent NT-sgRNA controls, and blue traces represent gene-specific sgRNA conditions. Histograms are shown for Total, CD15^+^, SOX2^+^, and CD15^+^/SOX2^+^ populations. Comparable shifts in fluorescence intensity across all subpopulations indicate consistent knockdown efficiency without subpopulation-specific differences. (H) Subpopulation analysis of CRISPRi-mediated KD of surface markers. KD efficiency for B2M and CD151 was quantified by flow cytometric analysis and expressed as percentage reduction relative to matched NT-sgRNA controls, calculated from isotype-corrected mean fluorescence intensity (MFI) values for each population. To evaluate whether KD efficiency differed among subpopulations, percentage reduction values were analysed using one-way repeated-measures ANOVA (RM-ANOVA) with subpopulation across three biological replicates (n=3). *P*-values were derived from the F-distribution corresponding to the ANOVA model. Data are presented as mean ± SD; non-significant comparisons are not indicated (*P* ≥ 0.05).

To assess CRISPRi activity in the engineered GBM cells, we performed sgRNA-mediated KD of the surface markers B2M and CD151. G7-CRISPRi cells transfected with sgRNAs targeting the promoter of these genes exhibited robust KD (>97%) at both the mRNA (Figure 1D) and protein levels (Figure 1E and 1F). Since the persistence of stem-like cells represents a major driver of radioresistance in GBM, we next evaluated gene KD efficiency in SOX2- and CD15-expressing stem-like subpopulations by flow cytometry analysis. Strong KD (>95%) of B2M and CD151 was also observed in the SOX2-positive, CD15-positive, and double-positive subsets, supporting the use of this CRISPRi-engineered GBM model for identifying genetic modifiers of radiation response (Figure 1G and 1H). Lastly, we observed that co-transfection of two sgRNAs targeting the same gene achieved greater KD efficiency than single sgRNA transfection (Supplementary Figure 1C). Therefore, this dual-sgRNA strategy was adopted for our ClonoScreen3D-CRISPRi screening workflow.

### Selection of candidate modifiers of radiation response to test ClonoScreen3D-CRISPRi screening workflow

We employed four distinct strategies to test whether the ClonoScreen3D-CRISPRi screening platform could be used to identify genes that modulate GBM radiosensitivity: **(i)** selection of genes significantly upregulated following ionizing radiation (IR) as identified by RNA-Seq^12^, **(ii)** inclusion of genes previously identified in our work as critical regulators of GBM growth and survival, specifically those involved in cholesterol homeostasis, a known driver of GBM progression; **(iii)** investigation of novel genes involved in the DNA damage response (DDR), including *CDK9, CDK8*, and *DSCC1*; and **(iv)** inclusion of genes strongly associated with radioresistance, such as ATM and PRKDC (DNA-PK), which are the targets of established radiosensitizing compounds in patient-derived GBM cell lines. These kinases are the targets of established radiosensitizing compounds and are central regulators of the DDR; ATM is primarily recruited to double-strand breaks (DSBs) where it orchestrates the signalling cascade for homologous recombination (HR) and cell cycle checkpoints, while PRKDC is essential for the non-homologous end joining (NHEJ) pathway.

To study the first cohort of potential targets we utilized analytical methods previously described^12^. This analysis leveraged RNA-Seq data from G7 cells cultured in 3D conditions and harvested four hours post-exposure to 5 Gy IR or sham treatment. We observed that IR significantly altered the expression profile of the non-canonical NF-κB pathway, including *NFKB2* and *RELB* transcripts (Figure 2A). These genes encode p100/p52 and RelB, respectively, integral components of the non-canonical signalling cascade^21^. Simultaneously, we observed a more modest induction of *NFKBIA* and *NFKBIE*, which encode the inhibitory regulators IκBα and IκBε, respectively. Given the roles of these proteins in sequestering canonical NF-κB dimers, these data indicate that IR-induced transcriptional changes preferentially favour upregulation and activation of the non-canonical NF-κB pathway over the canonical signalling axis (Figure 2A).

**Figure 2.**
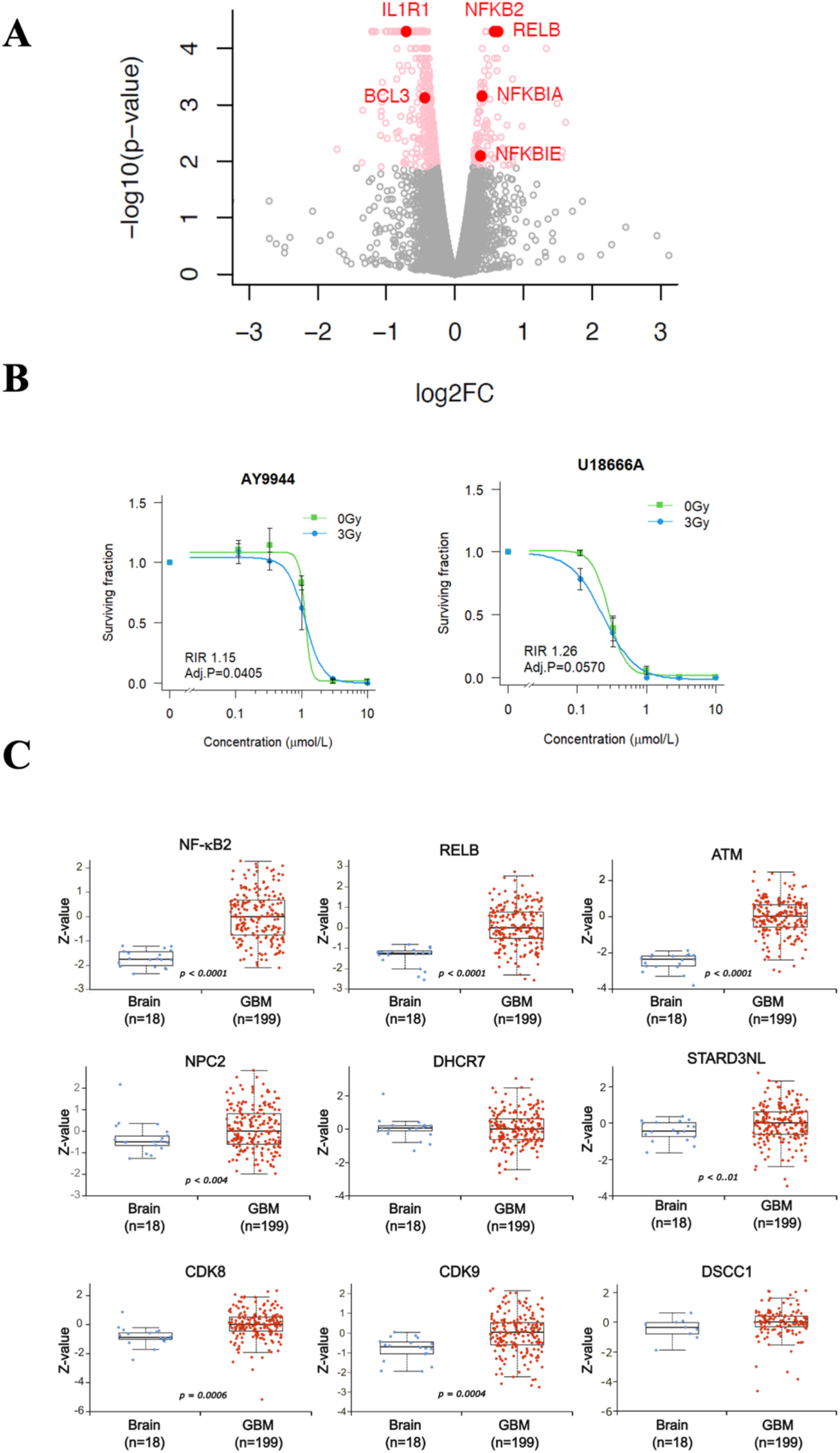
Selection of candidate genes to test ClonoScreen3D-CRISPRi screening workflow. (A) RNA-Seq-based identification of radiation driven changes in gene expression in the patient-derived G7 cell GBM cells. Volcano plot of RNA-Seq analysis of G7 GBM cells exposed to radiation (5 Gy vs untreated control (0 Gy)) and changes in gene expression determined by RNA-Seq analysis, with a focus on genes involved in the non-canonical NF-κB pathway, observing a 1.49, 1.54, 1.31 and 1.29 log_2_fold change in *NFKB2, RELB, NFKBIA* and *NFKBIE*, respectively. (B) Clonogenic survival of G7 cells following treatment with AY9944 (left panel) or U18666A (right panel) or vehicle two hours prior to IR (3 Gy) calculated from automated colony counts. The radiation interaction ratio (RIR) is calculated by comparison of areas-under-the-curve of the control (0 Gy) and irradiated (3 Gy) samples, following log-transformation of concentration. The surviving fraction of irradiated samples was computed using the plating efficiency of the vehicle + 3 Gy control, normalizing for the effect of radiation alone. Data shown as mean ± standard deviation (SD), *n*=3. EC_50_(mmol/L) with 95% confidence interval calculated by fitting of a 4-parameter dose response curve. EC_50_values compared by testing means of ratios. (C) Total protein expression pattern of candidate genes in glioblastoma CPTAC database. Boxplots generated using UALCAN showing total protein expression of NFKB2, RELB, ATM, NPC2, DHCR7, STARD3NL, CDK8, CDK9 and DSCC1. Z-score was calculated for each protein, defined as the number of standard deviations from the median across all samples. These normalized values were utilized to compare protein abundance in GBM tissues relative to healthy brain controls, with the resulting expression profiles.

For the second cohort, we focused on gene candidates that disrupt cholesterol metabolic pathways, specifically biosynthesis, represented by 7-dehydrocholesterol reductase (*DHCR7*) and 24-dehydrocholesterol reductase (*DHCR24*)^22,23^; and endosomal trafficking, represented by the Niemann-Pick Type C2 (*NPC2*) and STARD3 N-terminal like protein (*STARD3NL)*. Alterations in cholesterol metabolism, including abnormal activation of the *de novo* synthesis pathway^24^, impaired uptake^25^, and defective trafficking^26^, are key mechanisms driving GBM proliferation and treatment resistance. We found that targeting DHCR7 with AY9944^27^, as well as the dual-action compound U18666A, targeting DHCR24 and the NPC1, demonstrated potent single agent efficacy. AY9944 showed an IC_50_ of 0.288 µM (95% CI, 0.232-0.344 µM), while U18666A showed an IC_50_ of 1.12 µM (95% CI, 0.78-2.58 µM) at 0 Gy, with no radiation interactions observed (Figure 2B).

To confirm the clinical relevance of our selected gene targets, we interrogated protein expression levels in GBM tumours relative to normal healthy brain tissue using the CPTAC database and the UALCAN integrated cancer data analysis platform^20^. This proteomic analysis confirmed that several targets identified in our strategies are expressed in GBM samples, and in several cases, highly enriched, in GBM tissue compared to healthy controls, including NF-kB2 (*P*<0.0001), RELB (*P* <0.0001), ATM (*P* <0.0001), NPC2 (*P* =0.04), CDK8 (*P* <0.0001), CDK9 (*P* <0.0001) (Figure. 2C) and NFKBIA and NFKBIE (*P* <0.0001; Supplementary Figure 2). This validation ensured that the candidate genes used for testing our ClonoScreen3D-CRISPRi platform are representative of the proteomic landscape observed in patient-derived tumour samples. By corroborating our findings with CPTAC data, we established a robust link between our experimental gene cohorts and the clinical pathology of GBM.

### Development and validation of the ClonoScreen3D–CRISPRi clonogenic screening platform

In order to utilize the clonogenic survival assay as a readout for the ClonoScreen3D-CRISPRi screen, we developed a new automated colony counting pipeline, conceptually aligned with our previous approach^12^, incorporating Cellpose-based image segmentation followed by CellProfiler analysis to improve colony detection and quantification in Alvetex plates. (Figure 3A, 3B). A strong correlation was observed between manual counting (counting by eye) and the automated colony counts generated by the developed pipeline (Supplementary Figure 3).

**Figure 3.**
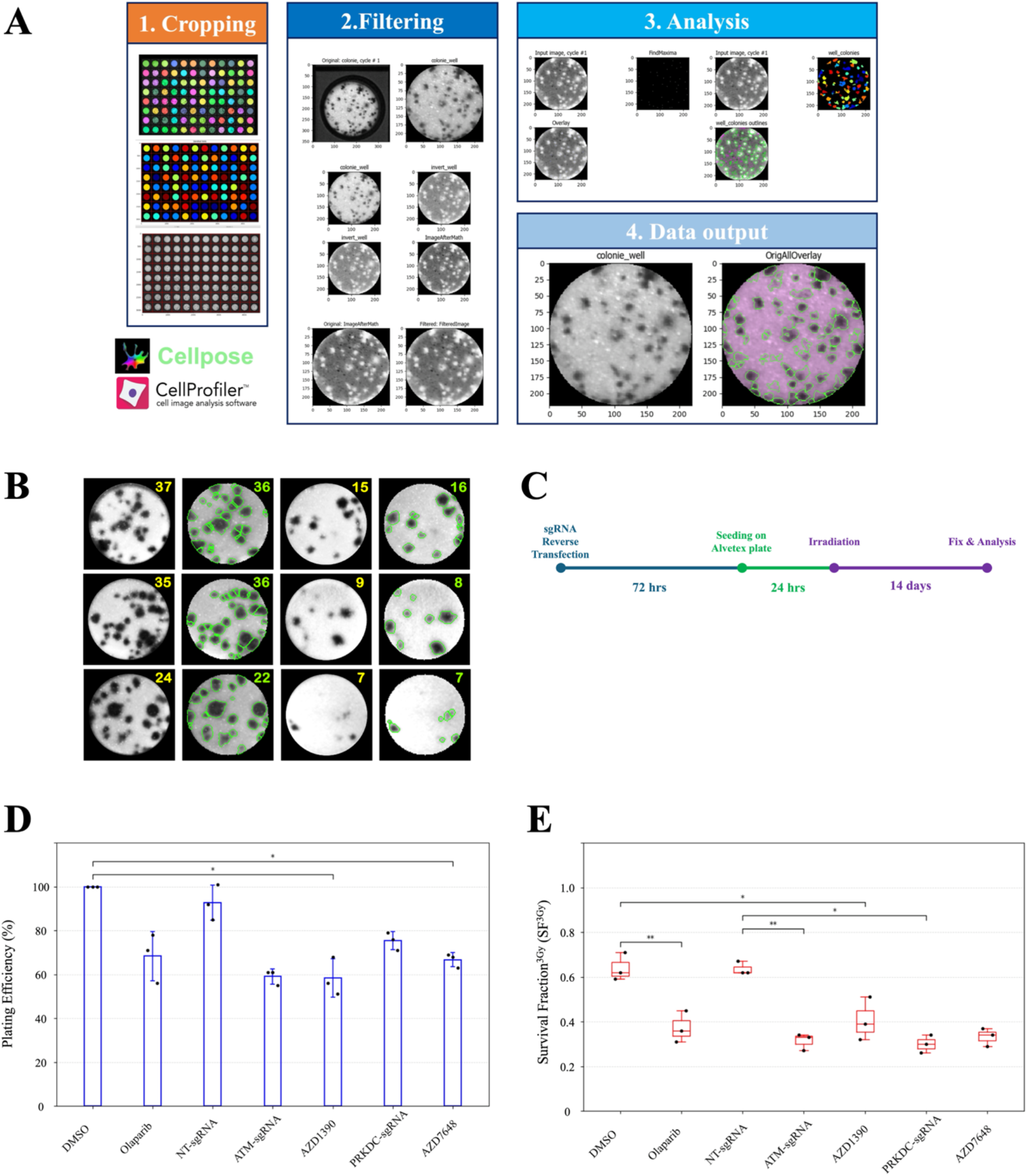
Development and validation of the ClonoScreen3D–CRISPRi clonogenic screening platform. (A) ClonoScreen3D–CRISPRi colony readout and image-based quantification pipeline. Representative images illustrate the semi-automated colony detection workflow. After fixation, raw images were first cropped using CellPose and subsequently processed for colony detection/counting using CellProfiler within our analysis pipeline. (B) Representative analysed images of colonies on Alvetex 96-well plates, demonstrating robust discrimination and numbers of individual colonies. Yellow numbers indicate manual counting, whereas green numbers represent automated counting. (C) Experimental timeline and workflow of the ClonoScreen3D–CRISPRi screening assay. CRISPRi-engineered GBM-Zim3 cells were reverse-transfected with sgRNAs and allowed 72 hours for transcriptional repression before seeding onto Alvetex plates. These plates were irradiated 24 hours after seeding, and maintained for 14 days to enable long-term clonogenic expansion. Colonies were then fixed and quantified, allowing assessment of KD- or compound-specific effects on post-irradiation clonogenic survival. (D) Functional validation of the ClonoScreen3D–CRISPRi platform using reference radiosensitizing perturbations. Plating efficiency (PE) under unirradiated conditions (0 Gy) was assessed following pharmacological inhibition of key DNA damage response components (ATM inhibitor AZD1390, 10 nM; DNA-PK inhibitor AZD7648, 10 nM; PARP inhibitor olaparib, 1 µM) or CRISPRi-mediated gene knockdown (KD). PE was calculated from colony counts and normalization to the non-targeting (NT) sgRNA control (for gene KD) or DMSO control (for compound treatments). Data are presented as mean ± SD from three independent biological replicates (n = 3). Statistical analysis was performed using repeated-measures one-way ANOVA followed by Dunnett’s multiple comparisons test against the corresponding control. \**P* < 0.05; non-significant comparisons are not shown. (E) Surviving fraction (SF) following irradiation (3 Gy) under the same pharmacological and CRISPRi perturbation conditions as in (D). SF was calculated from colony counts and normalization to the corresponding 0 Gy condition within each biological replicate. Data are presented as independent biological replicates (n = 3). Statistical analysis was performed using one-way ANOVA followed by Dunnett’s multiple comparisons test against the appropriate control. \**P* < 0.05, \*\**P* < 0.01; not significant comparisons are not shown.

We tested our ClonoScreen3D-CRISPRi workflow (Figure 3C) by performing KD of *PRKDC* and *ATM*, prior to radiation treatment, and compared the effects of these KDs with those of AZD1390 and AZD7648, which inhibit the kinase activity of ATM and DNA-PK respectively, and with the PARP inhibitor olaparib (Figure 3D and 3E). In the absence of irradiation, all three small molecule inhibitors impacted plating efficiency (PE) with both AZD1390 and AZD7648 significantly reducing PE compared with the DMSO control. Both gene KDs also impacted PE, but these effects were not significant compared with the non-targeting (NT)-sgRNA (Figure 3D). The surviving fraction (SF) following 3 Gy irradiation was reduced to 0.299 ± 0.043 with KD of PRKDC, closely resembling the effect of the DNA-PK inhibitor AZD7648 (0.334 ± 0.042) (Figure 3E). Similarly, KD of *ATM* decreased SF to 0.315 ± 0.039, comparable to pharmacological inhibition with AZD1390 (0.406 ± 0.094). These effects were similar to those seen with the PARP inhibitor olaparib (SF=0.370 ± 0.070; mean ± SD; Figure 3E). Collectively, these results validate our screening workflow as a method to identify modifiers of radiation response in GBM cells. Based on consideration of both its strong radiation interaction effect and its modest impact on PE, *PRKDC* KD was also included in subsequent ClonoScreen3D-CRISPRi screens, as benchmarks for potent radiosensitization.

### A mini ClonoScreen3D-CRISPRi screen identifies NF-κB non-canonical pathway genes as potent modifiers of radiation response

The nine genes identified in our candidate gene analysis were screened using the ClonoScreen3D-CRISPRi workflow in G7-Zim3 and GBML20-Zim3 cells. Each screen was carried out in triplicates, with good correlation of colony numbers evident in both cell lines (Supplementary Figure 4B). KD of four genes (*CDK8, NPC2, STARD3NL*, and *DHCR7*) was associated with significant reductions in PE in the absence of irradiation in both cell lines (Figure 4A). Specifically, in G7-Zim3 cells, PE at 0 Gy decreased to 20.18% ± 1.82% (*CDK8*), 37.19% ± 1.40% (*NPC2*), 60.76% ± 3.14% (*STARD3NL*), and 35.48% ± 2.02% (*DHCR7*). In GBML20-Zim3 cells, corresponding values were 20.56% ± 1.80% (*CDK8*), 38.62% ± 2.80% (*NPC2*), 51.08% ± 1.58% (*STARD3NL*), and 37.97% ± 4.71% (*DHCR7*) (mean ± SD).

**Figure 4.**
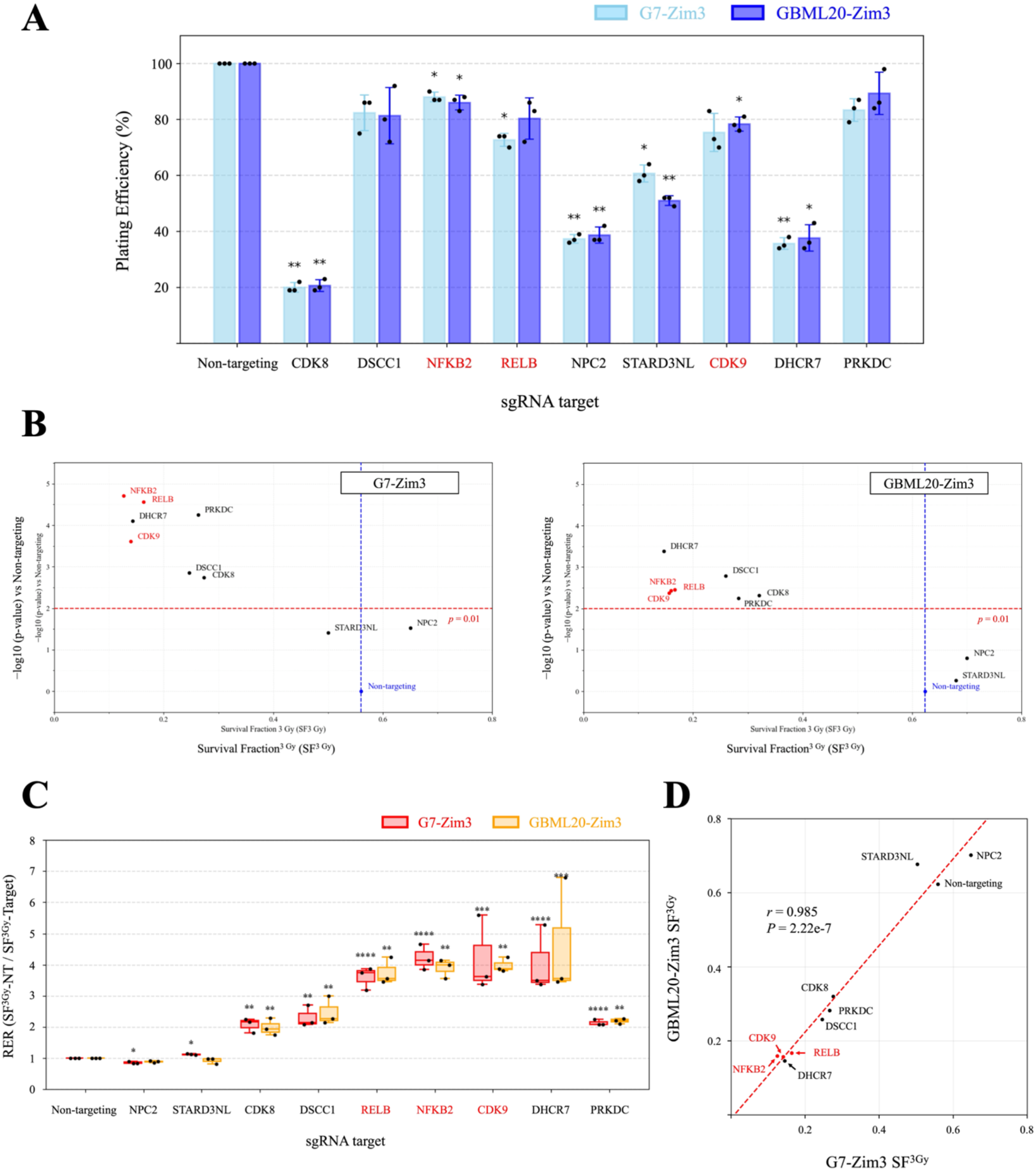
CRISPRi-based clonogenic radiosensitization screening in GBM cell lines. (A) Plating efficiency for each KD group was calculated by normalising colony numbers to the average colony count of the NT-sgRNA control group, which was defined as 100%. (B) Screening results are shown as scatter plots for the two CRISPRi GBM cell lines, G7-Zim3 (upper panel) and GBML20-Zim3 (lower panel), analysed using an identical experimental and analytical framework, with *P*-values compared to the NT-sgRNA control group. The surviving fraction (SF) at 3 Gy was calculated for each sgRNA condition by normalising the colony number obtained after irradiation (3 Gy) to the corresponding colony number under non-irradiated conditions (0 Gy) within the same sgRNA group. SF values were derived from three independent biological replicates (n=3). Statistical significance between each gene-specific knockdown and the NT-sgRNA control was assessed using one-way ANOVA followed by Dunnett’s multiple comparisons test. Adjusted P-values derived from the ANOVA model were obtained for each comparison and are presented as −log10(P). (C) Screening results are shown as a Radiation enhancement ratio (RER) for two CRISPRi GBM cell lines, G7-Zim3 and GBML20-Zim3. RER was calculated by as the ratio of SF^3Gy^ of NT-sgRNA to SF^3Gy^ of Target-sgRNA. RER value were derived from three independent biological replicates (n=3) are presented as mean ± SD. Statistical significance between each target KD and the NT-sgRNA control was assessed using one-way ANOVA followed by Dunnett’s multiple comparisons test. Adjusted P-values derived from the ANOVA model were obtained for each comparison and are presented as −log10(P). \**P* < 0.05, \*\**P* < 0.01 ***P < 0.001, ****P < 0.0001; not significant, ns, not shown). (D) Correlation of CRISPRi screening results. Scatter plot comparing SF following 3 Gy irradiation in G7-Zim3 and GBML20-Zim3 cells for all screened genes. The dashed line represents the least-squares linear regression fit. The significance of Pearson’s correlation coefficient (r) was assessed using a two-tailed t-test based on the t-distribution with n-2 degrees of freedom.

The ClonoScreen3D–CRISPRi screen revealed that KD of all candidate genes examined, with the exception of *STARD3NL* and *NPC2*, significantly altered the SF following 3 Gy irradiation in both cell lines (all *P* < 0.01) (Figure 4B and Supplementary Figure 4A). In G7-Zim3 cells, the most pronounced radiation interaction was observed following KD of *NFKB2* (SF = 0.125 ± 0.026), *CDK9* (0.141 ± 0.033), *DHCR7* (0.145 ± 0.031), and *RELB* (0.164 ± 0.024). A similar pattern was also observed in GBML20-Zim3 cells where KD of *DHCR7* (SF = 0.146 ± 0.039), *CDK9* (0.157 ± 0.002), *NFKB2* (0.159 ± 0.013) and *RELB* (0.167 ± 0.013) produced the strongest reductions in clonogenic survival.

Notably, KD of *CDK9, DHCR, NFKB2* and *RELB* had more potent effects on clonogenicity following irradiation than KD of known enhancer of radiosensitivity *PRKDC* (Figure 4B). The radiation enhancement ratio (RER) plot (Figure 4C), which reflects the extent to which gene knockdown enhances the radiation response, further indicated that KD of these genes produced the strongest radiosensitising effects, consistent with the screening results. These effects were consistent across both *MGMT* promoter methylated (G7-Zim3) and unmethylated (GBML20-Zim3) cell lines examined (Figure 4D). Among these genes, *CDK9, NFKB2* and *RELB*, were selected for further investigation to validate the impact of their KD on the radiosensitivity. Given the radiation independent cytotoxicity associated with KD of *DHCR7* (Figure 4A), this gene was not pursued for further investigation despite its potent radiation interaction effect on SF.

### KD of NF-κB non-canonical pathway genes elicits radiosensitization of GBM cells

We confirmed mRNA KD of *CDK9, NFKB2* and *RELB* by qPCR analysis of cells 72 hrs post sgRNA transfection before seeding onto Alvetex plates. Expression of all three genes was reduced by more than 60% in both G7-Zim3 and GBML20-Zim3 cells (Figure 5A). To determine the impact of individual gene KD on radiosensitivity, full radiation dose–response assays were performed in both cell lines (Figure 5B). Survival curves were generated across the radiation dose range and sensitizer enhancement ratios (SER; Figure 5B) calculated. These showed that KD of *NFKB2, RELB*, and *CDK9* increased radiosensitivity relative to NT-sgRNA–transfected controls: in G7-Zim3 cells, KD of *NFKB2, RELB*, and *CDK9* yielded SER values of 1.63 (95% CI 1.53–1.72, *P* = 6.39 × 10^−5^), 1.48 (95% CI 1.39–1.58, P = 6.38 × 10^−5^), and 1.39 (95% CI 1.30–1.48, P = 0.00147), respectively. Similarly, in GBML20-Zim3 cells, KD of *NFKB2, RELB*, and *CDK9* yielded SER values of 1.70 (95% CI 1.64–1.76, *P* = 7.81 × 10^−6^), 1.58 (95% CI 1.45–1.73, *P* = 7.25 × 10^−6^), and 1.49 (95% CI 1.41–1.57, *P* = 4.68 × 10^−7^), respectively. For all three genes, statistical analysis of SER values confirmed significant radiosensitization in both cell lines.

**Figure 5.**
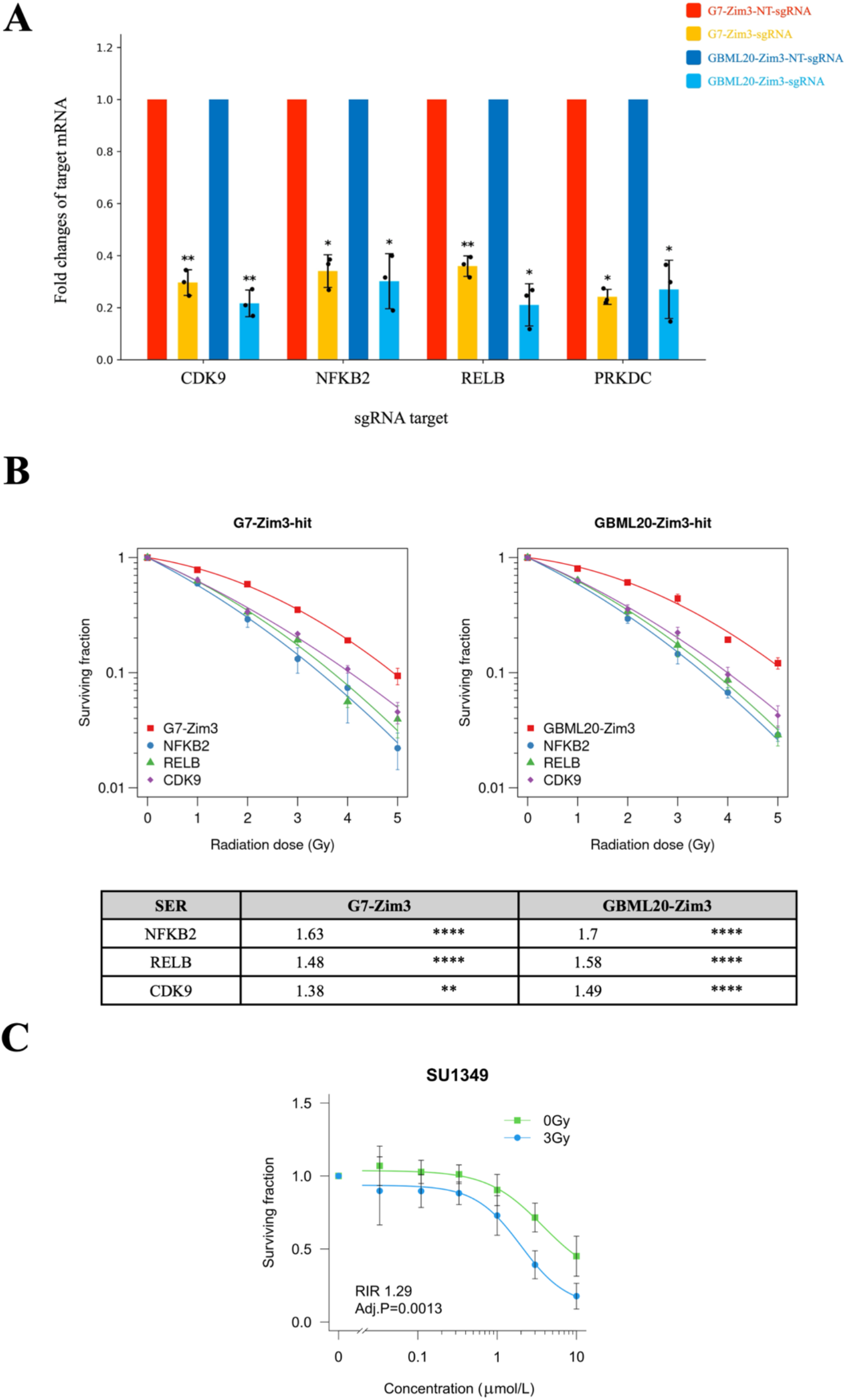
Validation of CRISPRi screening hits by genetic and pharmacological perturbation. (A) Relative mRNA expression levels of selected screening hits (*CDK9, NFKB2, RELB*, and *PRKDC*) were quantified by qPCR following sgRNA-mediated CRISPRi KD in G7-Zim3 and GBML20-Zim3 cells. Transcript levels were normalized to NT-sgRNA controls using the ΔΔCt method, confirming efficient and comparable knockdown across both GBM cell lines. Statistical significance was assessed using paired two-tailed Student’s t-tests on ΔCt values across biological replicates (n=3). Data are presented as mean ± SD, \**P* < 0.05, \*\**P* < 0.01; ; not significant comparisons are not indicated. (B) Validation of screening hits across a full radiation dose range. SF was assessed over increasing doses of ionising radiation in G7-Zim3 and GBML20-Zim3 cells following CRISPRi-mediated KD of selected hits. Radiation survival curves demonstrate consistent modulation of long-term clonogenic survival across the dose range, validating screening-identified effects beyond the single-dose (3 Gy) screening condition. Data fitted using the linear quadratic model. (C) Pharmacological validation of NF-κB–associated hits using pathway inhibition. Dose response analysis of clonogenic survival of G7 cells following treatment with SU1349 or vehicle two hours prior to IR (3 Gy) calculated from automated colony counts. The radiation interaction ratio (RIR) is calculated by comparison of areas-under-the-curve of the control (0 Gy) and irradiated (3 Gy) samples, following log-transformation of concentration. The surviving fraction of irradiated samples was computed using the plating efficiency of the vehicle + 3 Gy control, normalizing for the effect of radiation alone. Data shown as mean ± standard deviation (SD), *n*=3. EC_50_(mmol/L) with 95% confidence interval calculated by fitting of a 4-parameter dose response curve. EC_50_values compared by testing means of ratios.

To further validate the radiosensitizing effects of inhibition of non-canonical NF-kB pathway genes *NFKB2* and *RELB*, and the transcriptional regulator *CDK9*, we evaluated pharmacological inhibition of these targets in combination with ionizing radiation. G7 cells were treated with the inhibitor SU1349^28^, which acts as a potent IKKα inhibitor but has additional inhibitory activity against CDK9. This compound exhibited a significant radiation interaction in G7 cells (RIR 1.29; *P* = 0.0013) (Figure 5C), demonstrating that chemical inhibition of these pathways phenocopies the genetic KD results and reinforcing their therapeutic potential for GBM. Taken together, these data show that *NFKB2, RELB* and *CDK9* are radiosensitising targets in GBM cells and demonstrate that ClonoScreen3D-CRISPRi is a suitable method to identify such targets.

## Discussion

In this work we have established ClonoScreen3D-CRISPRi as a novel platform for identifying genetic modifiers of radioresistance in GBM. By combining CRISPRi-mediated gene KD with 3D clonogenic survival assays in patient-derived GBM cells, this platform addresses two key limitations of prior genetic screens in *in-vitro* human models: the use of 2D cultures that fail to recapitulate GBM complexity, and reliance on short-term viability readouts that do not capture long-term clonogenic capacity. The latter is a critical determinant of tumour recurrence following radiotherapy and the gold-standard endpoint for identification of modifiers of radioresistance.

Previous CRISPR-based screens in GBM have identified genetic drivers of temozolomide (TMZ) resistance and radiation response but were predominantly conducted in 2D monolayer cultures using short-term viability as the primary readout^7,9,10^. The CRISPRi-based radiation modifier screen by Liu *et al*. (2020) used clonogenic endpoints in glioma but focused on long non-coding RNAs and was performed in 2D^11^ while Lin *et al*. (2025) used CRISPRoff for epigenetic silencing in GBM, demonstrating chemo-sensitization but not radiation response^8^. A previous study performed genome-wide CRISPR knockout screens on a bioprinted 3D model of GBM and on GSCs cultured as spheres, however it only investigates gene essentiality and not radiation response^29^.

ClonoScreen3D-CRISPRi advances on these approaches by integrating gene silencing with 3D clonogenic readouts in patient-derived GBM lines with characterized *MGMT* status, increasing clinically relevance. Furthermore, the workflow is designed to accommodate larger focused libraries, expanding the scope for identifying novel radiosensitizing targets. Most importantly, our data demonstrate that efficient CRISPRi-mediated KD was achieved in stem-like subpopulations of GBM cells. Although these populations of GBM are widely recognized as central drivers of tumour recurrence and therapeutic resistance, defining functionally relevant genetic vulnerabilities within them has remained challenging^30,31^. Assessing gene function in stem-like populations provides a method to uncover radiosensitizing targets that have been insufficiently explored in the context of clonogenicity.

We found that seven out of nine candidate genes, which were selected by transcriptomics analysis, pharmacological inhibitor studies and a survey of the literature, significantly altered clonogenic SF following irradiation in the two GBM cell lines examined. Of note, silencing of *NFKB2, RELB*, or *CDK9* markedly suppressed clonogenic survival following 3 Gy irradiation. Interestingly, these KDs did not induce significant cytotoxicity in the absence of radiation, suggesting that expression of these gene is selectively required for post-irradiation recovery and maintenance of clonogenic potential rather than basal survival. The potency of our identified targets is striking. The SER values for KD of *NFKB2* (1.70), *RELB* (1.58), and *CDK9* (1.45) significantly exceeded the SER of 1.20 observed for the PARP inhibitor olaparib^12^. Furthermore, silencing these targets surpassed the effects of silencing *ATM* and *PRKDC*, which are often used as benchmarks for potent radiosensitization.

*NFKB2* encodes the central structural hub of the non-canonical NF-κB pathway, and *RELB* is its principal transcriptional effector. Through the regulated proteolytic processing of p100 into p52, *NFKB2* enables RELB-dependent transcriptional programs essential for stabilizing the mesenchymal (MES) state ^32^. The reduction in clonogenic survival following *NFKB2* or *RELB* KD is consistent with a role for the non-canonical NF-κB pathway in post-irradiation recovery. Traditionally, the canonical pathway (RelA/p65) is thought to handle immediate DNA damage responses and anti-apoptotic signalling, while the non-canonical pathway (p52/RelB) facilitates long-term cellular adaptation^33,34^. However, our findings do not conform to this strict division, rather suggesting a model whereby non-canonical NF-kB signalling contributes to radioprotective mechanisms in GBM.

*CDK9*, the catalytic subunit of the positive transcription elongation factor b (P-TEFb) complex, regulates RNA polymerase II pause release and is selectively involved the transcriptional response to cellular stress ^35^. The radiosensitization observed upon *CDK9* KD is likely to reflect a failure to execute the non-canonical NF-κB-driven MES program at the level of transcriptional elongation. Therefore, we hypothesise that CDK9 and IKKα-mediated non-canonical NF-κB signalling operate within the same regulatory axis to promote post-irradiation survival, though further mechanistic studies are needed to confirm this.

Beyond transcriptional regulation, the platform identified a dependency on cholesterol homeostasis: KD of *NPC2* (a lysosomal transporter) and *DHCR7* (the terminal enzyme in the Kandutsch-Russell pathway) exerted potent cytotoxic activity even without radiation, consistent with the known “cholesterol addiction” of GBM cells ^36^. In addition, a significant radiation interaction was observed following KD of *DHCR7*, underscoring the need to investigate DHCR7 further as a potential radiosensitization target in GBM. Our identification of these distal pathway components as survival-critical bottlenecks adds to the growing evidence for cholesterol metabolism as a therapeutic vulnerability in GBM ^37–39^.

Taken together, these results demonstrate that ClonoScreen3D-CRISPRi is an effective and scalable platform for identifying genetic modifiers of radioresistance in GBM. The use of 3D clonogenic endpoints in patient-derived stem-like cells provides a biologically relevant context that is better positioned to predict translational outcomes than conventional 2D viability screens. Future application of ClonoScreen3D-CRISPRi with larger CRISPRi libraries has the potential to uncover novel modifiers of radioresistance and to help prioritize targets for pharmacological development in GBM.

## Acknowledgements

This work was funded by NC3Rs [G122436]. We thank the NIHR Cambridge BRC Cell Phenotyping Hub for flow cytometry analysis. We thank members of the Target Discovery Team and the Chalmers lab for helpful discussions and support.

## Author contributions

N.M., E.B. and N. G.-R. conceived the work and designed experiments, S.L. executed the experiments and analysed the data, A.H. developed the automated analysis pipeline for the clonogenic assays, J.L. contributed to target validation experimets, C.Z.L. and S.M. contributed to the generation of the CRISPRi lines, S.A. advised and trained on the irradiation treatments, S.M., A.P., M.R.J and A.C. contributed to candidate genes selection, target validation and provided input into the manuscript. S.L., E.B. and N. G.-R. wrote the manuscript with input from all other authors.

## Materials and Correspondence

All correspondence and requests for materials should be addressed to Erica Bello.

