## Supplementary Information for "ClonoScreen3D-CRISPRi Uncovers Genetic Modifiers of Radiation Response in Glioblastoma"

**Consisting of**

Supplementary Tables 1-4

Supplementary Figures 1-4

Supplementary figure legends

**Supplementary Tables**

**Supplementary Table S1. sgRNA**

| Gene ID | sgRNA 1 | sgRNA 2 |
| --- | --- | --- |
| Non-Targeting | ACGACTAGTTAGGCGTGTA | N/A |
| B2M | CGCGAGCACAGCTAAGGCCA | N/A |
| CD151 | CCCGGACTCGGACGCGTGGT | N/A |
| ATM | TTTGAACCGGAAGCGGGAGT | GTAGGTAGCTGCGTGGCTAA |
| PRKDC | AGCGGGACTCGGCGGCAUGG | CAGCAGGGAGCAACGCACAC |
| CDK8 | GGGCAGAGCCGCACAACAGC | CAGGGAGCCCGCGGGGACAA |
| DSCC1 | CGCGGCGGGACTCCTAGACC | TTCCCAAGCAGCCGGAAGGT |
| NFKB2 | ATCCGGTGGAGAGCGAGATC | TCCCCGGCCAAGCCCAACTC |
| RELB | GTCCACCAGACCGTGCCTCC | GGGTCGGACGAGCGGCGCAA |
| NPC2 | GATAACGAAGTTCCAAGCTC | GGATAACGAAGTTCCAAGCT |
| STARD3NL | GTGACCGGGACCCAGTCCAG | CGCCCGCACCCAGCAGCCCC |
| CDK9 | GCGGCGGCAGCAGCGACTGG | CGGAGGGGCCTGGAGTGCGG |
| DHCR7 | GCCGCCTACCCTCTAGCCAG | GCGCGCGCAAGCGAGGCCAG |

**Supplementary Table S2. Compounds**

| Name | Manufacturer | Cat no. |
| --- | --- | --- |
| AZD2281 (olaparib) | Selleckchem | S1060 |
| AZD1390 | Selleckchem | S8680 |
| AZD7648 | Selleckchem | S8843 |

**Supplementary Table S3. Antibody**

| Antibody | Fluorophore | Species | Manufacturer | Cat no. |
| --- | --- | --- | --- | --- |
| SOX2 | VB423 | Mouse | Miltenyi Biotec | 130-131-185 |
| CD15 | FITC | Mouse | BioLegend | 323003 |
| B2M | APC | Mouse | BioLegend | 395711 |
| CD151 | APC | Mouse | BioLegend | 350405 |
| Isotype IgG1_k_ | BV421 | Mouse | BioLegend | 400157 |
| Isotype IgG1_k_ | FITC | Mouse | BioLegend | 400109 |
| Isotype IgG1_k_ | APC | Mouse | BioLegend | 400121 |

**Supplementary Table S4. qPCR primer**

| Gene ID | Forward | Reverse |
| --- | --- | --- |
| *ACTB* | ATTGGCAATGAGCGGTTC | GGATGCCACAGGACTCCAT |
| *B2M* | GAGTATGCCTGCCGTGTG | AATCCAAATGCGGCATCT |
| *CD151* | CATCGCTGGTATCCTCG | CTCGCTGCCCACAAAG |
| *ATM* | TCCGAGTGCAGTGACAGTGA | AGGATCTCGAATCAGGCGCT |
| *PRKDC* | AAGGCGGCTTACCTGAGTGA | CCTCACGAAGGCCCGCTTTA |
| *CDK8* | AAAGTGAAGCTGAGCAGCGA | TTCCCATCTTTCCTCTTGGCT |
| *DSCC1* | TGGTTGTAAAACTCCGGACCA | GCTTCTTTAACTTGGGTCTACGTC |
| *NFKB2* | TCCTCAAGATGGGGAGACTTCA | GTTTGGGGACAGTGAACAAGTG |
| *RELB* | CCTCTGGGCCGTCCGTC | ACTCGTCGATGATCTCTGTGC |
| *NPC2* | GCCTGATGGTTGTAAGAGTGG | TCACCCCCAGATAGACTTACGA |
| *STARD3NL* | GTTGTCCGCTGGACTGGG | TAACGGGTGGTTCCTGGC |
| *CDK9* | AAAGCAGTACGACTCGGTGG | CAGCACCTTCTTCAGAGCCA |
| *DHCR7* | GAGGTGTGCGCAGGACTTTA | TGGCTTTGGGAATGTTGGGT |

**Supplementary Figures**

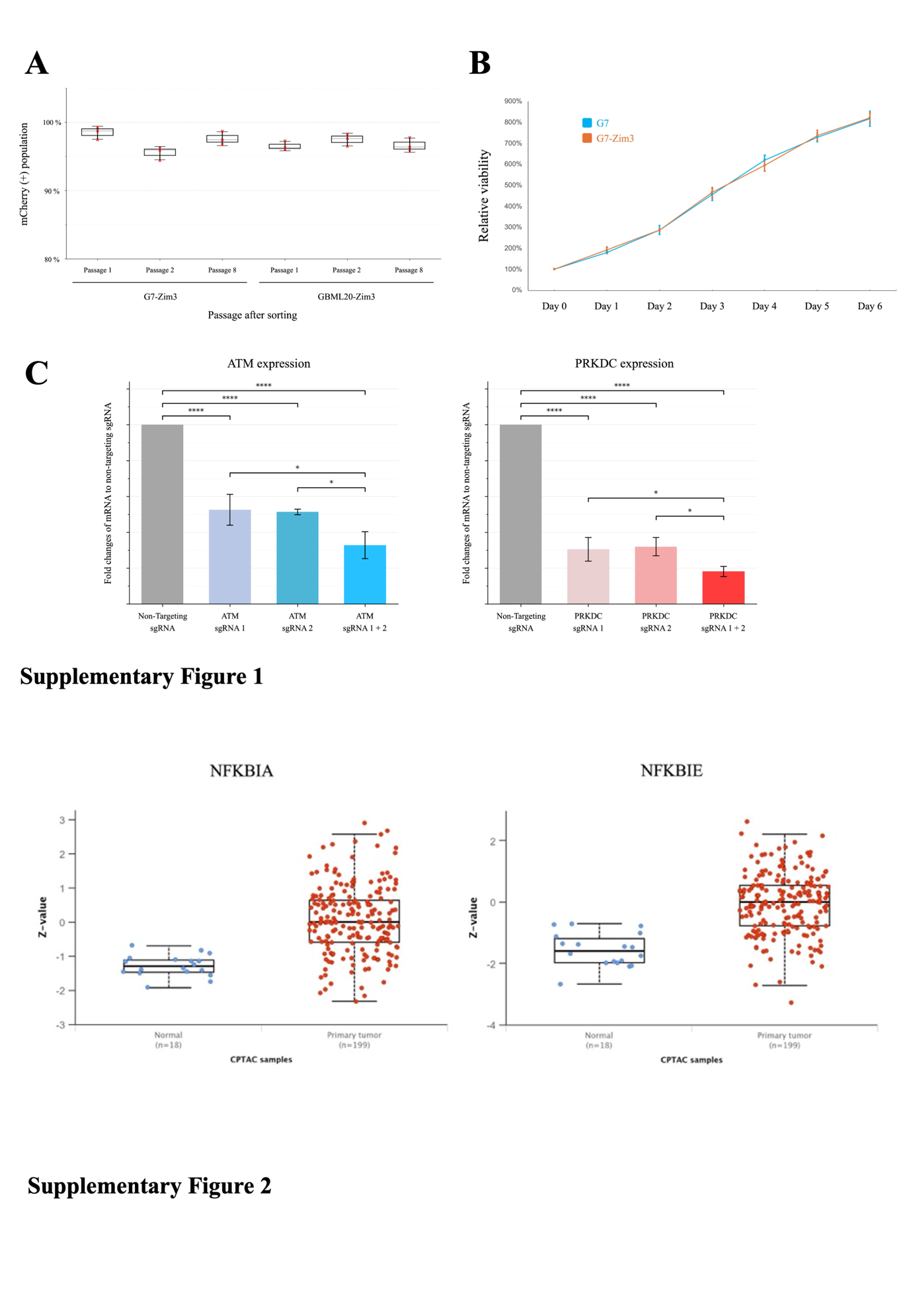

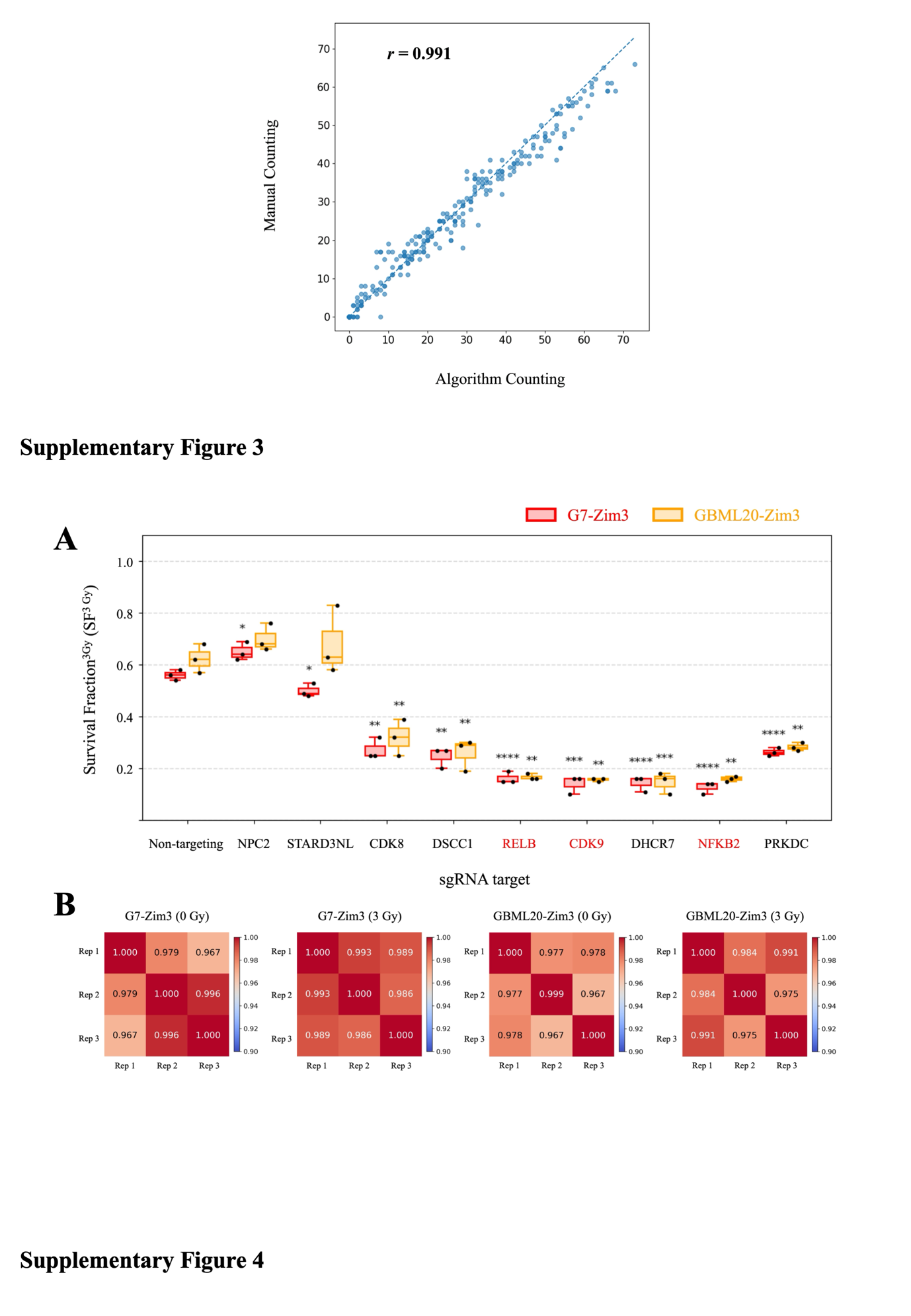

**Supplementary figure legends**

**Supplementary Figure 1.**

(A) Stability of CRISPRi-marked cell populations (mCherry^+^) after cell sorting using flow cytometry. The percentage of mCherry^+^ cells was quantified by flow cytometry at indicated passages, after mCherry^+^ sorting of G7-Zim3 and GBML20-Zim3 cell lines, respectively, demonstrating sustained expression of the CRISPRi construct over serial passaging.

(B) Growth analysis of parental G7 and CRISPRi-engineered G7-Zim3 cells. Cell growth was measured using the RealTime-Glo MT Cell Viability Assay. A total of 1,000 cells per well were seeded, and luminescence was recorded at baseline (day 0) and every 24 hours for 6 days. Luminescence values were normalized to the corresponding day 0 signal. Data represent mean ± SD from triplicate wells.

(C) qPCR validation of CRISPRi-mediated knockdown using single and dual sgRNAs. Gene knockdown efficiency was assessed by qPCR following reverse transfection of sgRNAs. Cells were harvested 96 hours post-transfection, and relative mRNA expression levels of *ATM* (left) and *PRKDC* (right) were quantified. Expression levels were normalised to non-targeting (NT) sgRNA control cells, defined as 1 (fold change = 1). KD efficiency was compared between single sgRNA and dual sgRNA (sgRNA1 + 2) conditions. Data represent mean ± SD from independent experiments. Statistical analysis was performed using one-way ANOVA followed by Tukey’s multiple comparisons test. Significance is indicated as *P < 0.05, **P < 0.01, ***P < 0.001, ****P < 0.0001.

**Supplementary Figure 2.**

Total protein expression pattern of NFKBIA and NFKBIE in glioblastoma CPTAC database. Boxplots generated using UALCAN showing total protein expression of NFKBIA and NFKBIE. The Z-score was calculated for each protein, defined as the number of standard deviations from the median across all samples.

**Supplementary Figure 3.**

Automated versus manual colony counting. Colony counts from four randomly selected plates (n = 384 wells) were analysed on a paired well-by-well basis. Pearson correlation analysis was performed to assess the linear relationship between automated and manual counting methods (r = 0.991). The dashed line represents the line of identity (y = x), indicating equal colony counts between the two methods.

**Supplementary Figure 4.**

(A) Box plots depict representative SF distributions under 3 Gy conditions for selected perturbations. The surviving fraction (SF) at 3 Gy for each sgRNA condition was determined by dividing the colony count following irradiation (3 Gy) by the matched non-irradiated (0 Gy) colony count from the same sgRNA group. SF values were obtained from three independent biological replicates (n=3), represented as closed dots in the box plot.

(B) Pearson correlation analysis of colony numbers obtained from three biological replicates under identical experimental conditions in G7-Zim3 and GBML20-Zim3 cells (0 Gy and 3 Gy). For each dataset, replicate-level vectors containing colony counts across all experimental conditions were correlated pairwise to generate 3 × 3 correlation matrices (Pearson’s r), and visualized as heatmaps. High concordance across replicates underscores the robustness and reliability of the ClonoScreen3D-CRISPRi as a screening platform.
